# The Influence of Magnetic Fields on Selected Physiological Parameters of Blood and Tissues in Mice

**DOI:** 10.1101/497990

**Authors:** Hani M. Abdelsalam, Mohammed Elywa

## Abstract

This study aimed to highlight the influence of exposure to different applied magnetic fields (MFs) on SOD, MDA and GSH levels in the liver, LDH and CPK activities in the muscle and γ-aminobutyric acid levels in the brain, as well as some haematological parameters. Adult male albino Swiss mice were divided into 5 equal groups (*n* = 6), the control group (untreated) and four exposure groups that were exposed to MFs of 20, 40, 60 and 80 Gauss for 5 min/day for 5 days.: Exposure to MFs induced significant decreases in total GSH levels and SOD activity but a significant increase in MDA levels in the liver. By contrast, SMF exposure significantly increased total LDH and total CPK activities in the muscle. The results revealed a significant increase in GABA levels in the brain, as well as decreases in haemoglobin, haematocrit, and red blood cell counts, in addition to platelet counts, after exposure to 20, 40, 60 and 80 Gauss MFs. After exposure to a 40 Gauss MF, the mice showed pathological changes in red blood cells, including changes to the outer membrane of the red blood cells (micronucleus and a serrated edge, with a mild incidence of echinocytes). In the group exposed to a 60 Gauss MF, examination of blood smears clearly showed changes in cell size, with the emergence of abnormal forms, including many areas with no red blood cells (rouleaux formation). With increasing intensity of exposure (80 Gauss), the red blood cells appeared completely different from their natural form and took the form of ovalocytes and bi-micronucleated erythrocytes, which appear in patients with anaemia. MF exposure caused different metabolic and haematological effects, which appeared to be related to the intensity of SMF exposure. The changes in the biochemical parameters of SMF-exposed mice probably reflect hepatic damage and anaemia caused by kidney failure. Further studies are needed to obtain a better understanding of the effects of MF on biological systems.

## Introduction

The planet is surrounded by magnetic fields (MFs) generated by the earth, solar storms, variations in the weather and everyday electrical events. Recently, scientists have discovered that external MFs can affect the body in both positive and negative ways, and such clinical observations have revealed new avenues of study. The ability of high static magnetic fields (SMFs) to provide higher resolution and more frequent spectral changes in magnetic resonance imaging (MRI) and spectroscopy (MRS) has increased. However, to assure patient safety, a full understanding of the effects of MFs on human physiology is required [16].

Indices related to red blood cells (RBCs), which can serve as markers of red blood cell function, provide a clear picture of the performance and efficiency of red blood cells, and haemoglobin concentrations can describe increases and reductions in the volume of red blood cells [7].

Exposure to SMFs induces metabolic and haematological changes that correlate with the length of exposure. Moreover, exposure to voltage reduces RBC function and metabolic activity; thus, it was proposed that increased toxicity in organs was an outcome of RBC failure [8].

Rapid and precise signal transmission among the majority of nerve cells (neurons) in the mammalian central nervous system (CNS) is mediated by two major modes of neurotransmission: excitation by glutamic acid and inhibition by γ-aminobutyric acid (GABA). GABA is generated from glutamic acid by the action of glutamic acid decarboxylase (GAD) [46]. [54] GABA and glutamate levels in the brain cortex, striatum, hippocampus, and hypothalamus were determined, and the results demonstrated the strong effect of GABA and glutamate levels on the hippocampus and hypothalamus. However, GABA receptor modulators did not induce a substantial effect on extremely low-frequency magnetic field (ELF-MF)-induced changes or increased levels of GABA at the tested dose.

In mammals, LDH occurs in two isomeric forms: the M subunit from skeletal muscle and the H subunit from heart muscle [25]. Increases in serum LDH may result from the release of LDH isoenzymes from both the muscle and the heart. Furthermore, the detection and quantification of LDH isozyme levels in the blood may reveal possible organ toxicity of SMFs. In SMF-exposed rats, physiological adaptation to hypoxia might alter the endocrine system [51].

Creatine kinase (CK) or creatine phospho-kinase (CPK) is assayed in blood tests as a marker of damage in CK-rich tissue, such as in patients with myocardial infarction (heart attack), rhabdomyolysis (severe muscle breakdown), muscular dystrophy, autoimmune myositis, and acute kidney injury **[35]**.

Radiation increases heart enzymes in animals and has been proposed to cause cardiovascular disease in these animals **[11]**. Additionally, **[3]** reported that male white mice exposed to phone calls 10 times per day, for 10 min each, every day for one month showed effects on creatine phosphokinase, LDH, AST and high-density lipoprotein, which reflect cardiac function; in addition, differences in pathological damage were observed in the hearts of the group exposed to mobile phone rays compared to those of the control group.

Although oxygen is required for many important aerobic cellular reactions, it may undergo electron transfer reactions, which generate highly reactive membrane-toxic intermediates, such as superoxide, hydrogen peroxide and the hydroxyl radical **[30]**. Numerous environmental factors may interfere with this phenomenon, including long-term exposure to ELF-MF **[26]**.

Many studies have revealed the effects of electromagnetic field (EMF) exposure on total antioxidant activity, including SOD, GPx, vitamin E and A concentrations, MDA (a product of polyunsaturated fatty acid peroxidation that is used as an indicator of oxidative stress in cells and tissues) and selenium concentrations in erythrocytes and the plasma **[27]**. MDA is a highly toxic molecule that has been implicated in a range of disease pathologies by producing oxidative damage in tissues **[21]**.

In previous studies performed by **[29]**, human and mouse fibroblasts did not exhibit alterations in SOD, GPx, or FELINE activity when MFs with flux densities of 0.49 T or 128 mT were applied. However, studies on the effects of SMFs on rats reported discrepant outcomes. **[28]** showed that the activity of GPx in the muscles and kidneys and the activity of SOD in the liver were increased by SMFs. In contrast to **[24]**, chronic application of EMFs to mice does not cause oxidative damage, as indicated by a lack of SOD, GSH-Px, and catalase induction.

The time of application of MFs influenced biomass and GSH concentrations in yeast **[55]**. [31] reported that there is a correlation between exposure to SMF and oxidative stress through a disturbed redox balance, which leads to physiological perturbances. GSH is an efficient ROS scavenger that is essential for GPx and glutathione S-transferase activities and is considered the first line of antioxidant defence **[36]**.

## Methods

### Experimental animals

Forty adult male albino Swiss mice (*Mus musculus)* weighing 25-30 g were used in the present study. The animals were maintained under normal conditions and given food and water *ad libitum*. The mice were classified into 5 equal groups as follows:

I- Control group: The animals were left untreated as normal controls.
II- 20 Gauss group: The animals were exposed to 20 Gauss MFs for 5 min/day for 5 days.
III- 40 Gauss group: The animals were exposed to 40 Gauss MFs for 5 min/day for 5 days.
IV- 60 Gauss group: The animals were exposed to 60 Gauss MFs for 5 min/day for 5 days.
V- 80 Gauss group: The animals were exposed to 80 Gauss MFs for 5 min/day for 5 days.

### Source of Animals

The animals were obtained from the Theodor Bilharz Research Institute in Cairo, Egypt, and all animal procedures were performed after approval from the Ethics Committee of the National Research Center (ECNRC) in Egypt and in accordance with recommendations for the proper care and use of laboratory animals. The mice were killed by cervical dislocation after light anaesthesia (ether).

## Experimental setup

The experimental setup is illustrated in Fig. 1.

**Figure 1.**
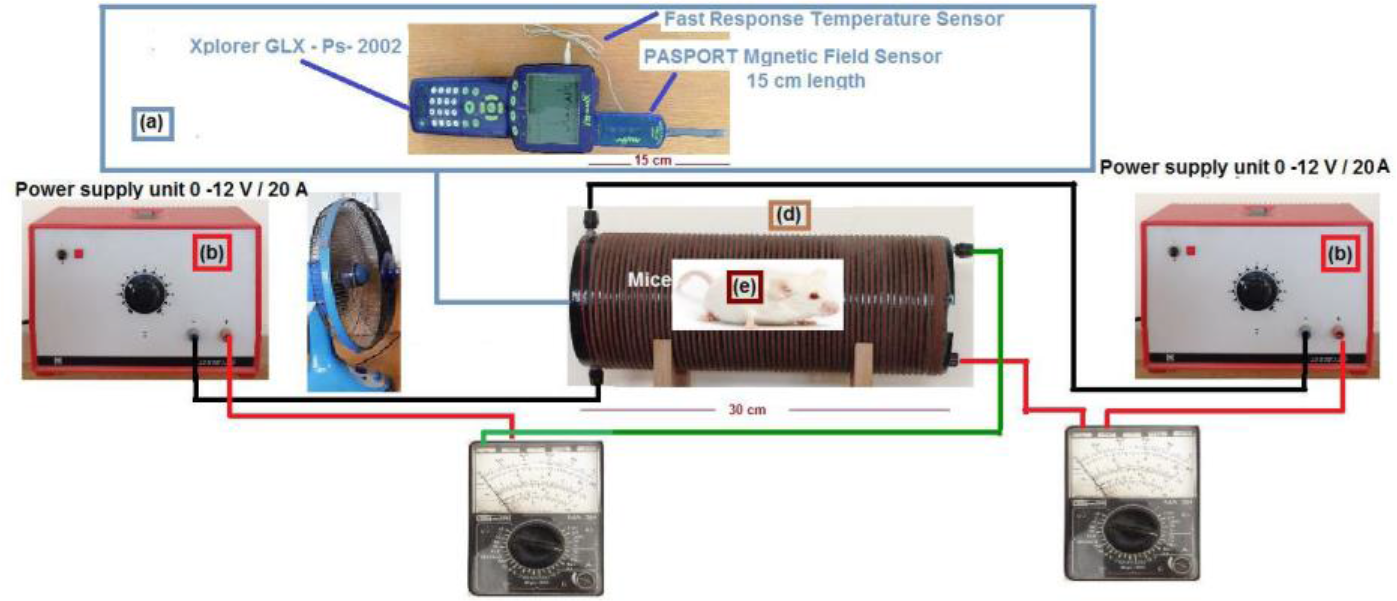
Schematic diagram of the SMF circuit. a) Xplorer GLX −PS-2002 with temperature and magnetic field probes used to measure the magnetic field intensity and the temperature simultaneously. b) a regulated d. c. voltage of low ripple. c) Multimeter to measure the d. c, current. d) double wrapped coil used to produce static magnetic field. e) a mice cage. And using a fan to cool the temperature rising. **This is designed by Co-author: Mohammed Elywa**

**Fig. (2):**
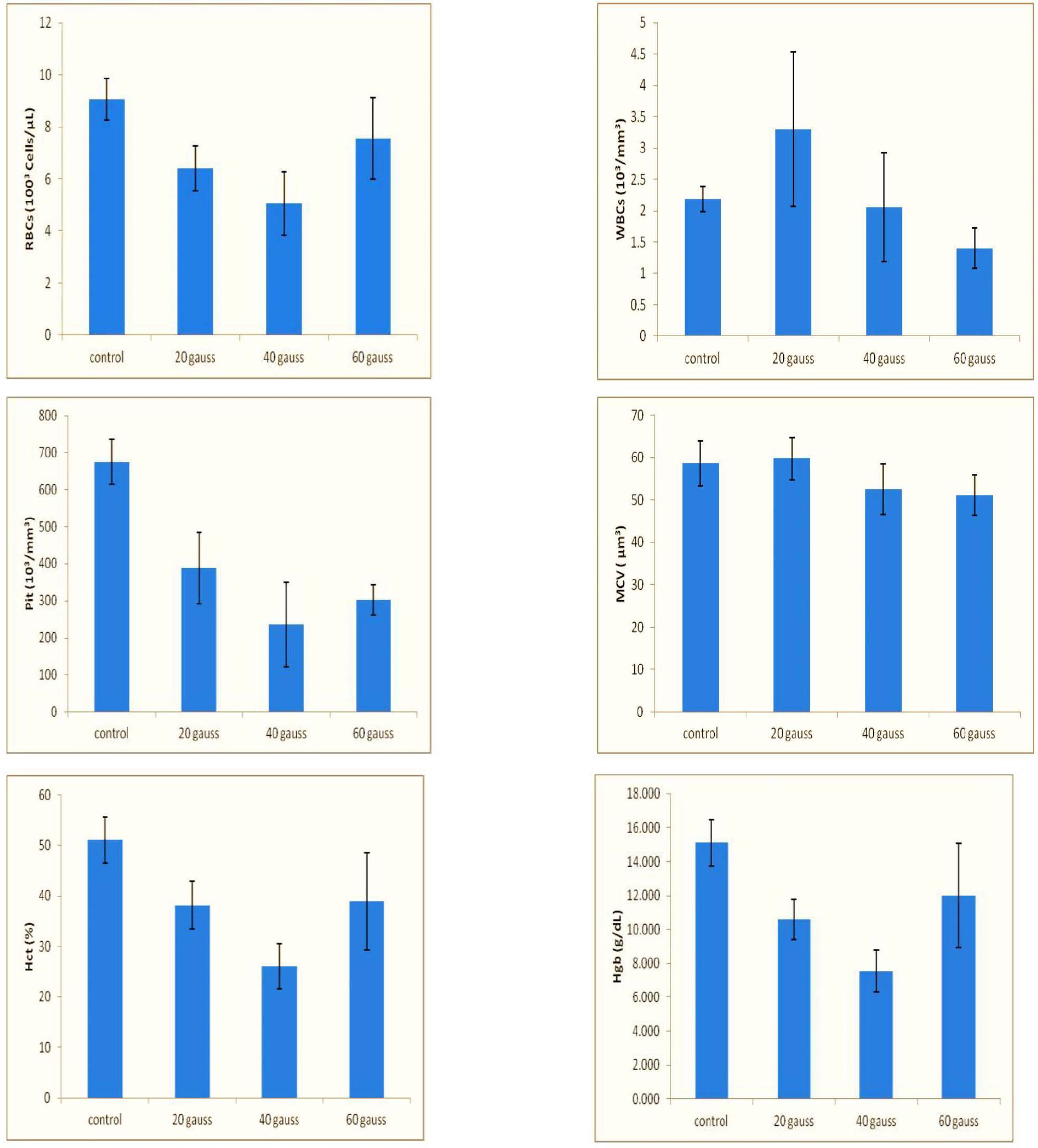
Haematological parameters of the mice exposed to 20, 40, 60 and 80 Gauss magnetic fields for 5 min/day for 5 days. Histopathological Results:

**Fig. (3):**
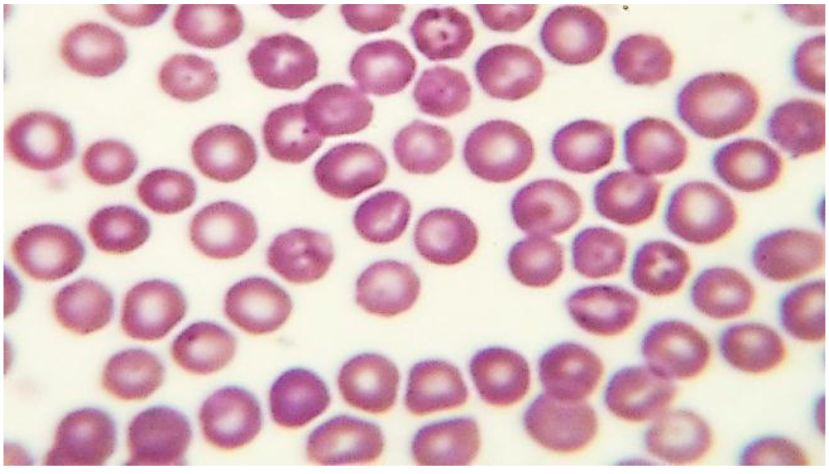
Representative photo of a Giemsa-stained blood smear from the control group showing normal disc-shaped biconcave erythrocytes (arrows) with a central pale area.

**Fig. (4):**
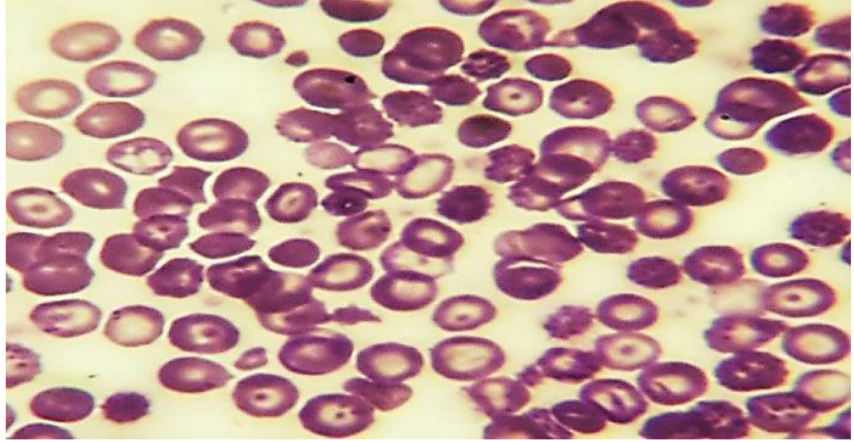
Representative photo of a Giemsa-stained blood smear from the 20 Gauss group showing a moderate incidence of echinocytes (black arrow), marked hypochromasia (red arrow), target cells (yellow arrows) and microcytes (blue arrows)

– A coil with a variable number of turns per unit length was placed on a stand for coils and tubes and connected to a high-current power supply.
– The axial B-probe was connected to the teslameter via the multicore cable, clamped with the stand rod from the probe apparatus, and aligned so that the Hall sensor (a) was located in the centre of the plastic body of the coil. **This is designed by Co-author: Mohammed Elywa**

## Physiological analysis

Haematological parameters were determined by using the haemocytometer method for RBC counts, WBC counts and platelet counts; the Wintrobe microhaematocrit method for PCV; and the Drabkin method for Hb determination, according to ***[22]***.

Lipid peroxidation was estimated by measuring the level of thiobarbituric acid reactive substances (TBARS) in tissues by the method of ***[44]***.

CK is also known as creatine phosphokinase (CPK) or phospho-creatine kinase. CK was estimated by the method of ***[48]***.

Superoxide dismutase was assayed using the method of **[20]**.

GSH concentrations were determined by the method of ***[43]***.

GABA concentrations were determined by the method of ***[57]***.

### Preparation of tissue homogenates

Small pieces of the brain, liver, and muscle were collected and rinsed in 10% buffered neutral formalin solution. The remainder of the tissue (0.5 g) was independently homogenized in 5 mL of cold phosphate-buffered saline (pH 7.4, 0.1 M) using a Universal Laboratory Aid homogenizer (MPW-309, Mechanika Precyzyjna, Warsaw, Poland), filtered and centrifuged (BOECO centrifuge C-28A, Hamburg, Germany) at 3000 rpm for 15 min at 4 °C. The supernatants containing cell suspensions were collected and stored at 20 °C until further use in bioassays ***[42]***.

### Statistical analysis

The results presented here are the mean ± SE of 6 mice in each group. The results were analysed using one-way analysis of variance [ANOVA], and the group means were compared using Duncan’s multiple range test [DMRT] using SPSS version 12 for Windows. The findings were considered statistically significant if *P*<0.05 ***[10]***.

## Results

As shown in Table (1), mice exposed to a 20 Gauss MF showed a non-significant increase in the brain GABA level, with a percentage difference of 3.26%. Mice exposed to 40, 60 and 80 Gauss MFs showed a significant increase in brain GABA levels, with percentage differences of 15.75%, 33.46% and 52.47%, respectively, compared to those of the control group.

**Table (1):** GABA concentration (μg/g of tissue) in the brains of the mice exposed to 20, 40, 60 and 80 Gauss magnetic fields for 5 min/day for 5 days.

| Groups | Treatments | GABA ($\mu\text{g/g}$ of tissue) | P value | Percentage difference (%) |
| --- | --- | --- | --- | --- |
| Gp. I | Control | $178.83 \pm 1.96^d$ | $>0.05$ | - |
| Gp. II | 20 Gauss | $184.67 \pm 1.85^d$ | | 3.26 |
| Gp. III | 40 Gauss | $207.00 \pm 4.72^c$ | $<0.05$ | 15.75 |
| Gp. IV | 60 Gauss | $238.67 \pm 6.35^b$ | | 33.46 |
| Gp. V | 80 Gauss | $272.67 \pm 6.39^a$ | | 52.47 |

As shown in Table (2), mice exposed to a 20 Gauss MF showed a non-significant decrease in the level of muscle CPK, with a percentage difference of −8.38%. Mice exposed to 40, 60 and 80 Gauss MFs showed a significant increase in the levels of muscle CPK, with percentage differences of 170.38%, 226.68%, and 267.57%, respectively, compared to those of the control group.

**Table (2):** CPK activities (μmol/min/mg of protein) in the muscles of the mice exposed to 20, 40, 60 and 80 Gauss magnetic fields for 5 min/day for 5 days.

| Groups | Treatments | CPK ($\mu\text{mol/min/mg}$ of protein) | value | Percentage difference (%) |
| --- | --- | --- | --- | --- |
| Gp. I | Control | $88.77 \pm 1.68^e$ | $>0.05$ | - |
| Gp. II | 20 Gauss | $81.33 \pm 2.59^e$ | | -8.38 |
| Gp. III | 40 Gauss | $240.00 \pm 7.75^c$ | $<0.05$ | 170.36 |
| Gp. IV | 60 Gauss | $290.00 \pm 20.81^{ab}$ | | 226.68 |
| Gp. V | 80 Gauss | $326.30 \pm 18.44^a$ | | 267.57 |

As shown in Table (3), mice exposed to a 20 Gauss MF showed a non-significant increase in the hepatic MDA level, with a percentage difference of 28.57%. Mice exposed to 40, 60 and 80 Gauss MFs showed a significant increase in hepatic MDA levels, with percentage differences of 192.85%, 350.00%, and 478.57%, respectively, compared to those of the control group.

**Table (3):**
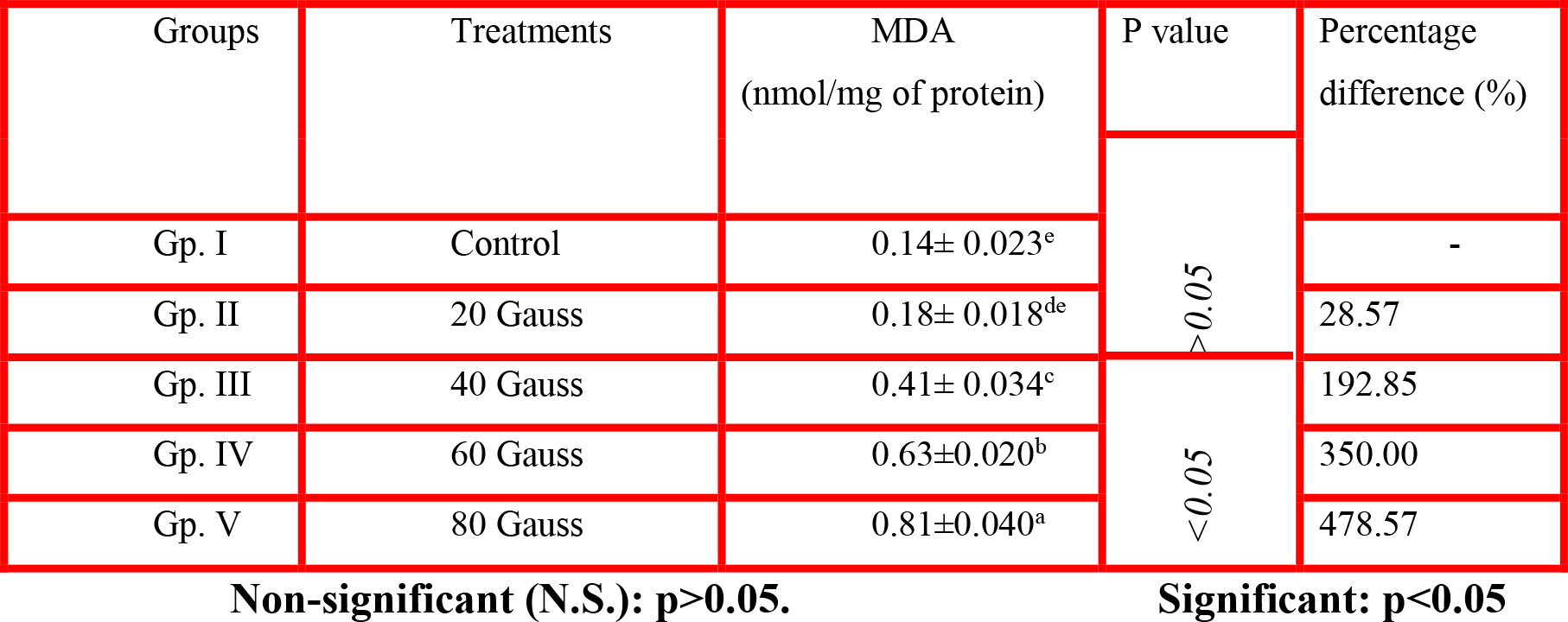
MDA levels (nmol/mg of protein) in the liver of the mice exposed to 20, 40, 60 and 80 Gauss magnetic fields for 5 min/day for 5 days.

As shown in Table (4), mice exposed to a 20 Gauss MF showed a non-significant decrease in hepatic SOD activity, with a percentage difference of −0.28%. Mice exposed to 40, 60 and 80 Gauss MFs showed a significant decrease in hepatic SOD activity, with percentage differences of −47.29%, −55.84% and −64.67%, respectively, compared to that of the control group.

**Table (4):** SOD activities (U/mg of protein) in the liver of the mice exposed to 20, 40, 60 and 80 Gauss magnetic fields for 5 min/day for 5 days.

| Groups | Treatments | SOD (U/mg of protein) | P value | Percentage<br>difference (%) |
| --- | --- | --- | --- | --- |
| Gp. I | Control | 3.51±0.044<sup>a</sup> | >0.05 | - |
| Gp. II | 20 Gauss | 3.50±0.061<sup>a</sup> |  | -0.28 |
| Gp. III | 40 Gauss | 1.85±0.031<sup>d</sup> | <0.05 | -47.29 |
| Gp. IV | 60 Gauss | 1.55±0.026<sup>e</sup> |  | -55.84 |
| Gp. V | 80 Gauss | 1.24±0.029<sup>f</sup> |  | -64.67 |

As shown in Table (5), mice exposed to 20 Gauss MFs showed a non-significant decrease in hepatic GSH content, with a percentage difference of −1.78%. Mice exposed to 40, 60 and 80 Gauss MFs showed a significant decrease in hepatic GSH content, with percentage differences of 66.07%, 83.92% and 92.85%, respectively, compared to that of the control group.

**Table (5):**
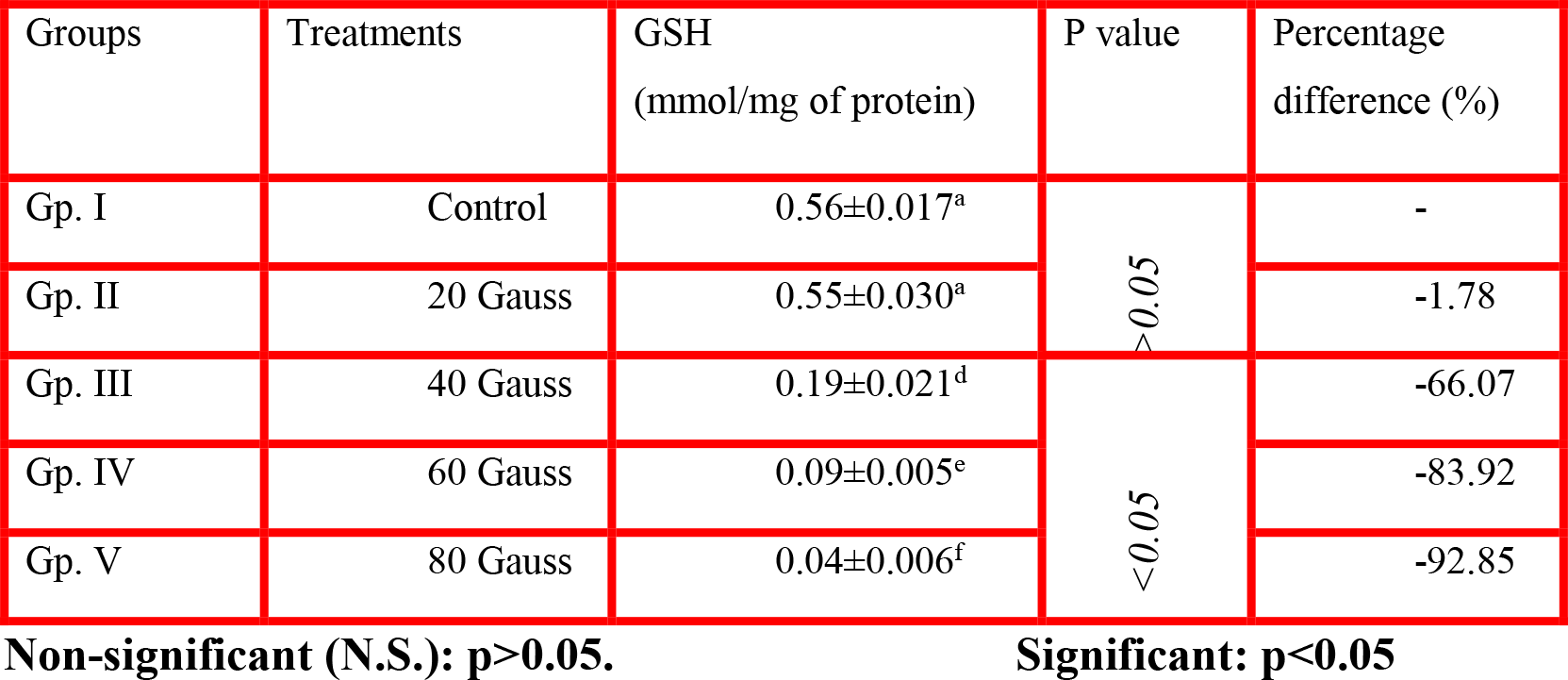
GSH levels (mmol/mg of protein) in the liver of the mice exposed to 20, 40, 60 and 80 Gauss magnetic fields for 5 min/day for 5 days.

| Groups | Treatments | GSH<br>(mmol/mg of protein) | P value | Percentage<br>difference (%) |
| --- | --- | --- | --- | --- |
| Gp. I | Control | 0.56±0.017<sup>a</sup> | >0.05 | - |
| Gp. II | 20 Gauss | 0.55±0.030<sup>a</sup> |  | -1.78 |
| Gp. III | 40 Gauss | 0.19±0.021<sup>d</sup> | <0.05 | -66.07 |
| Gp. IV | 60 Gauss | 0.09±0.005<sup>e</sup> |  | -83.92 |
| Gp. V | 80 Gauss | 0.04±0.006<sup>f</sup> |  | -92.85 |

As shown in Table (6), mice exposed to a 20 Gauss MF showed a non-significant decrease in the level of muscle LDH, with a percentage difference of -0.62%. Mice exposed to 40, 60 and 80 Gauss MFs showed a significant increase in the level of muscle LDH, with percentage differences of 76.14%, 123.38%, and 157.83%, respectively, compared to that of the control group.

**Table (6):** LDH activities (μmol/min/mg of protein) in the muscles of the mice exposed to 20, 40, 60 and 80 Gauss magnetic fields for 5 min/day for 5 days.

| Groups | Treatments | LDH<br>(μmol/min/mg of protein) | P value | Percentage<br>difference (%) |
| --- | --- | --- | --- | --- |
| Gp. I | Control | 33.87±0.81<sup>e</sup> | >0.05 | - |
| Gp. II | 20 Gauss | 33.66±2.96<sup>e</sup> |  | -0.62 |
| Gp. III | 40 Gauss | 59.66±5.48<sup>c</sup> | <0.05 | 76.14 |
| Gp. IV | 60 Gauss | 75.66±1.76<sup>b</sup> |  | 123.38 |
| Gp. V | 80 Gauss | 87.33±3.84<sup>a</sup> |  | 157.83 |

**Table (7):** Significant differences were observed in most of the haematological blood parameters (Hb, RBCs, Hct, Plt, MCV and WBCs) between the control and the exposed groups of mice after exposure to 20, 40, 60 and 80 Gauss MFs. MF exposure produced a more significant decrease (*p<0.001*) in Hb, RBCs, Hct and Plt after 20 and 40 Gauss MF exposure and a significant decrease (p <0.05) in Hb, RBCs and Hct after 60 Gauss MF exposure, but the decrease in Plt was more highly significant. In contrast, mice exposed to an 80 Gauss MF showed significant decreases in both Hb and Hct and non-significant decreases in both RBCs and Plt in comparison to those of the control group. While the WBC count was significantly increased after exposure to 20 Gauss and 80 Gauss MFs, with percentage differences of 50.86% and 47.62%, respectively, this value decreased non-significantly after exposure to a 40 Gauss MF and was more significant after exposure to a 60 Gauss MF, with percentage differences of −6.29% and −36.00%, respectively, in comparison to the control group. On the other hand, MCV showed a non-significant increase after exposure to a 20 Gauss MF and a decrease after exposure to both 40 and 80 Gauss MFs, whereas a significant decrease was observed after exposure to a 60 Gauss MF, with percentage differences of −12.77%.

**Table (7):** Haematological parameters of the mice exposed to 20, 40, 60 and 80 Gauss magnetic fields for 5 min/day for 5 days.

|  | Control | 20 Gauss |  |  | 40 Gauss |  |  | 60 Gauss |  |  | 80 Gauss |  |  |
| --- | --- | --- | --- | --- | --- | --- | --- | --- | --- | --- | --- | --- | --- |
|  | Mean±SE | Mean±SE | P value | % diff. | Mean±SE | P value | % diff. | Mean±SE | P value | % diff. | Mean±SE | P-value | % diff. |
| **Hb** | 15.10±0.33 | 10.60±0.19 | <0.001 | -29.82 | 7.55±0.08 | <0.001 | -50.01 | 12.00±0.22 | <0.05 | -20.55 | 12.40±0.30 | <0.05 | -17.93 |
| **RBCs** | 9.06±0.19 | 6.40±0.07 | <0.001 | -29.38 | 5.05±0.02 | <0.001 | -44.28 | 7.55±0.11 | <0.05 | -16.69 | 8.44±0.24 | >0.05 | -6.90 |
| **Hct** | 51.04±3.02 | 38.20±2.1 | <0.001 | -25.16 | 26.07±1.03 | <0.001 | -48.92 | 38.95±2.8 | <0.05 | -23.69 | 43.75±2.49 | <0.05 | -14.29 |
| **MCV** | 58.67±3.87 | 59.79±4.09 | >0.05 | 1.91 | 52.58±3.25 | >0.05 | -10.38 | 51.18±2.12 | <0.05 | -12.77 | 54.01±3.02 | >0.05 | -7.94 |
| **Plt** | 675.00<br>±39.6 | 388.00<br>±22.4 | <0.001 | -42.52 | 235.50±18.4 | <0.001 | -65.11 | 302.00±20.1 | <0.001 | -55.26 | 664.58±40.6 | >0.05 | -1.54 |
| **WBCs** | 2.19±0.26 | 3.30±0.34 | <0.05 | 50.86 | 2.05±1.08 | >0.05 | -6.29 | 1.40±0.03 | <0.001 | -36.00 | 3.23±0.27 | <0.05 | 47.62 |

*P values refer to significant differences compared to the control group.*
*Non-significant (N.S.): $p>0.05$ .*
*Significant: $p<0.05$ .*
*Highly significant: $p<0.01$ .*
*More highly significant: $p<0.001$*

## Discussion

Our studies showed a substantial increase in GABA levels in the brains of mice exposed to 40, 60 and 80 Gauss MFs for 5 min/day for 5 days. This observation is consistent with **[45]**, which reported that the exposure of adult rats to electromagnetic radiation (EMR) for 1 h daily for four months induced substantial increases in glutamate, aspartate, GABA and glycine levels but a strong decrease in glutamine levels.

An increase in cerebral metabolism was observed immediately after electromagnetic irradiation over some exposed regions of the scalp, and this effect was interpreted as a consequence of altered activity induced by EMF exposure **[33]**. This increase in cerebral metabolism may result in an increase in the rate of cerebral glucose utilization, which represents the main energy source in the brain **[56]**. Glucose is metabolized into amino acid neurotransmitters in neurons and glia **[17]**. Thus, the rapid increase in brain GABA levels after exposure of the mice to 40, 60 and 80 Gauss MFs for 5 min/day for 5 days in the present study may be due to an increase in brain metabolism induced by MFs.

The increase in aspartate and glycine levels might induce a state of hyperexcitability. However, an increase in brain amino acid levels is also a mechanism by which the brain ameliorates the state of hyperexcitability. In addition, might this increase in amino acid levels occur at the expense of less abundant amino acids? It has been urged that fifty of the endogenous amino acids are generated from other amino acids **[13]**.

Concerning the significant elevation of muscle CPK and LDH activities in mice exposed to 40, 60 and 80 Gauss MFs, our results are consistent with those reported in **[1]**, which demonstrated that rats exposed to an EMF for approximately 18 h every day for 3 months showed elevated CPK and LDH levels. **[52]** reported that hypoxia provoked a rapid loss of cellular ATP, followed by tissue functional and structural alterations, as revealed by an increase in LDH release. Thus, the increase in LDH activity following exposure to SMFs could indicate an adaptation that promotes anaerobic production of ATP in SMF-exposed rats **[2]**. Thus, the increase in LDH activity in the plasma following SMF application could be associated with severe muscle damage.

The present results showed that mice exposed to 40, 60 and 80 Gauss MFs showed a significant increase in hepatic MDA levels (0.41± 0.034, 0.63±0.020 and 0.81±0.04, respectively), with percentage differences of 192.85%, 350.00% and 478.57%, respectively. The present data showed a progressive increase in MDA levels with an increasing intensity of exposure. This relationship was accepted by **[9]**, who showed an elevation in MDA levels in the liver and excretory organs, indicating oxidative stress in response to SMFs (128 mT, one h/day for thirty consecutive days).

Our results are consistent with those of **[53]**, who reported that mice exposed to low-dose EMF exhibit potential harmful effects caused by enhanced radical production, as indicated by significant increases in the levels of antioxidant products, such as MDA and NO. However, MDA represents a breakdown of the major redox chain, leading to oxidation of polyunsaturated fatty acids, and thus serves as a reliable marker of oxidative stress mediated by oxidation **[58]**. These effects are associated with a higher rate of oxidative metabolic activity and a higher concentration of readily oxidizable membrane polyunsaturated fatty acids **[39]**. EMFs prolong the life of free radicals and can act as a promoter or co-promoter of cancer **[19]**.

Regarding SOD and GSH, the results of this study revealed that mice exposed to 40, 60 and 80 Gauss MFs showed significant decreases in hepatic SOD and GSH activities. These findings are consistent with **[53]**, who reported that immature female mice exposed to EMF for 4 h daily for 2 months, at 3 mT and 50 Hz, exhibited significant decreases in SOD, GPx and TAC activities compared to those of the control group. In this regard, **[4]** showed that exposure to EMF decreases GPx activity in ovarian tissue and uterine tissue in rats. GPx is an antioxidant enzyme that uses glutathione to reduce lipid hydroperoxides and hydrogen peroxide to reduce oxidative damage **[41]**. A decrease in GPx activity suggests the excess use of glutathione and reflects increasing levels of tissue MDA and oxidative damage. The decreased GPx activity and increased MDA levels increase in the tissues of the groups exposed to MFs in this study reveal an increase in lipid peroxidation. SOD is also an important antioxidant system enzyme that decomposes superoxide anion radicals to H_2_O_2_. In this way, the toxicity of superoxide is eliminated, and free radicals are not generated by superoxide **[49]**. These alterations may affect many processes within the cell, as free radical formation induces changes in enzyme activity, gene expression and membrane structure **[34]**.

The present results showed a significant reduction in the values of haemoglobin (Hb), RBCs, haematocrit (Hct) and platelets (Plt) in most mice exposed to 20, 40, 60 and 80 Gauss MFs in comparison to those in mice from the control groups, while the WBC count was increased in the 20 and 80 Gauss groups but decreased in the 40 and 60 Gauss groups in comparison to that in the control group. This finding is consistent with research that demonstrated that there are substantial decreases in the amounts of several factors in the blood, including most Hb, the Hct, RBCs, MCV, MCH, and MCHC **[5]**. Additionally, those authors observed a significant increase in the average WBC count, as well as the proportion of lymphocytes, and this increase was accompanied by cases of anaemia, such as macrocytic anaemia. This increase is also evidence of bleeding, which occurs as a protective mechanism after exposure to radiation and increased temperature **[18]**. These results are consistent with those of **[40]**, who stated that living near electrical relay networks leads to leukaemia. Additionally, **[38]** proved a relationship between exposure to electromagnetic waves and blood cancer (leukaemia). On the other hand, **[12]** showed that rats exposed to SMFs had increased WBCs in comparison to those of control rats. In the current study, blood smears were collected at the end of the experiment after exposing the animals to SMFs and investigated to study changes in the shape and size of RBCs and record consequences of defects or changes in these cells; these changes are of particular importance because they can be used to estimate the extent of environmental stress to which the animals were exposed. In this experiment, mouse blood samples were collected after exposure to 20, 40, 60 and 80 Gauss SMFs separately for 5 min/day for 5 days and then stained with Giemsa stain. The RBCs of mice exposed to 20 Gauss MFs showed a moderate incidence of echinocytes with a central pale area, marked hypochromasia and microcytic cells. These results were observed by **[23]**, who found that these forms or different abnormal forms of RBCs, termed erythrocyte poikilocytosis, were present in the smears, as were variations in the sizes of red blood cells, which were called anisocytosis. Some cells were a pale colour with an increased rate of colour fading in the swabs from mice that were exposed to electromagnetic waves; this observation has been attributed to a reduced amount of haemoglobin inside RBCs **[23]**.

Increased exposure to 40 Gauss MFs appears to have a clear influence on the blood swabs, with a mild incidence of echinocytes and micronuclei in the centre. Additionally, the sizes of RBCs differed from normal size, with clear areas between RBCs **(Fig. 5)**, which suggests that the lysis of many cells leads to a substantial reduction in the number of RBCs, as shown in **Table (7)**, which reported a percentage difference of -44.28%. The present result showed that SMFs have a stronger effect on the blood, where a reduced intensity of colour inside the RBCs appears due to a reduced concentration of haemoglobin (hypochromic); this finding was also confirmed in **Table (7)**, where a decrease in the average concentration of haemoglobin inside RBCs was observed after exposure to 40 Gauss MFs, with a percentage difference of −50.01%. It was noted that the mice exposed to 60 Gauss exhibited varying RBC sizes (anisocytosis). Microscopic observation revealed that the aggregation of RBCs and rouleaux formation occurred in the plasma, and rouleaux were oriented in the direction in which their long axes were perpendicular to the magnetic field **[15]**. This type of behaviour occurs in swabs from patients with anaemia and haemolytic anaemia **[32]**.

**Fig. (5):**
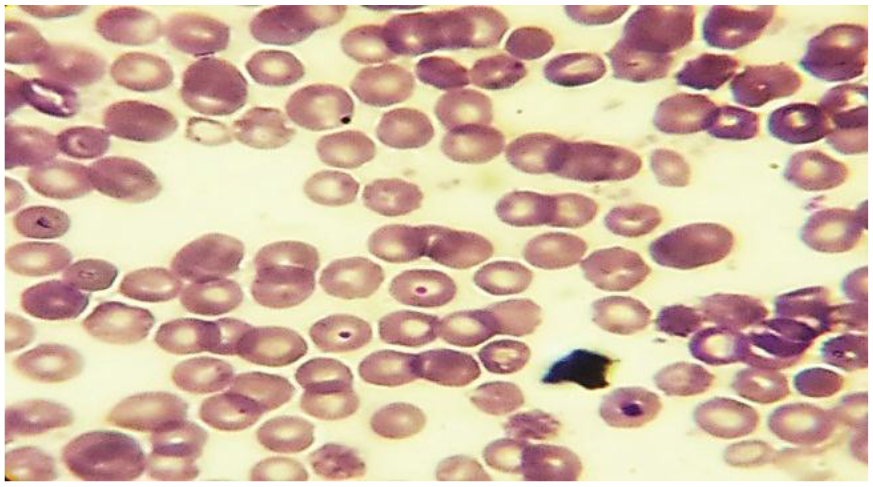
Representative photo of a Giemsa-stained blood smear from the 40 Gauss group showing a mild
3 incidence of echinocytes (black arrow) and micronuclei (blue arrows)

**Fig. (6):**
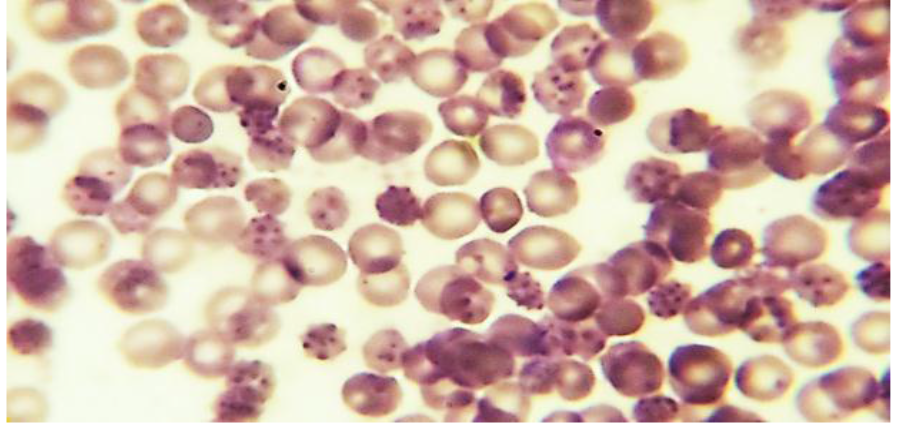
Representative photo of a Giemsa-stained blood smear from the 60 Gauss group showing a moderate incidence of echinocytes and the formation of a rouleaux that may lead to a clot.

**Fig. (7):**
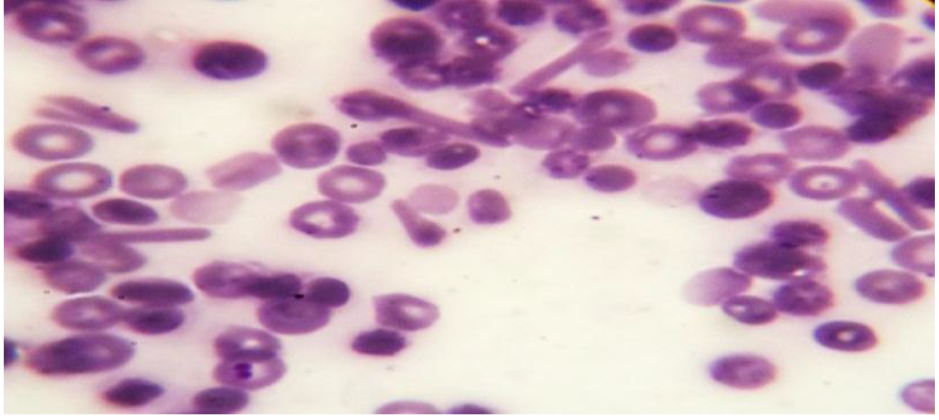
Representative photo of a Giemsa-stained blood smear from the 80 Gauss group showing a mild incidence of echinocytes (black arrow), bi-micronucleated erythrocytes (red arrow), and ovalocytes (blue arrows).

In this part of the study, the mice were exposed to 80 Gauss MFs 5 min/day for 5 days. Many pathological changes in RBCs were observed after exposure. RBCs with serrated walls were observed, with different numbers of humps, and other cells showed a structure surrounding the RBCs from the outside. These RBCs appear with a serrated edge and are called burr cells or called echinocyte erythrocytes (mild incidence of echinocytes). Bi-micronucleated erythrocytes and ovalocyte cells were also observed [37], confirming that these abnormal forms of RBCs were due to the presence of anaemia or anaemia caused by kidney failure.

In the current study, changes and a clear imbalance in the forms and sizes of RBCs and their contents occurred after exposure to SMFs; this effect may be attributed to a malfunction **[59]** and a clear change in the form of the membrane that surrounds the newly formed RBCs, which leads to an imbalance in the function of the membrane due to electromagnetic waves. **[23]** mentioned that the cell membrane of the globule is what gives it the flexibility necessary to pass through capillaries and control the exchange of elements inside and outside the RBCs; external stimuli can lead to a change in the composition and shape of the membrane, thereby hindering the establishment of this natural membrane cycle, which leads to the emergence of some serious diseases. This observation is consistent with the possibility outlined in **[6]** that the exposure to electromagnetic waves from mobile phones in humans leads to damage and has a clear influence on cell membranes, especially the membranes of RBCs, causing an imbalance in blood enzymes.

### Conclusion

MF exposure caused different metabolic and haematological effects, which appeared to be related to the intensity of SMF exposure. The changes in the biochemical parameters of SMF-exposed mice probably reflected hepatic damage and anaemia caused by kidney failure. Further studies are needed to better understand the effects of MFs on biological systems.

## Abbreviations

MF: Magnetic field
SMF: Static magnetic field
EMF: Electromagnetic field
SOD: Superoxide dismutase
GSH: Glutathione
MDA: Malondialdehyde
GABA: gamma-Aminobutyric acid
CPK: Creatine phosphokinase
LDH: Lactate dehydrogenase
RBC: Red blood cell
Hb: Haemoglobin
MCV: Mean corpuscular volume
WBC: White blood cell
Plt: Platelets

## Declarations

### Ethical approval and consent to participate

The animals were obtained from the Theodor Bilharz Research Institute in Cairo, Egypt, and all animal procedures were performed after approval from the Ethics Committee of the National Research Center (ECNRC). I agree to provide related documentation.

### Consent for publication

All authors consented to the publication of this manuscript.

### Availability of data materials

The data are publicly available in the Supplementary File.

### Author contributions

HA carried out the physiological and histological studies, participated in the sequence alignment and drafted the manuscript. ME participated in the experiments presented in the article. **All authors read and approved the final version of the manuscript**.

## Acknowledgements

I would like to acknowledge my parents, who instilled in me the value of education and the rewards and opportunities it can generate. In particular, my father supplied me with enthusiasm, support and creative insight. His critical reading of the manuscript helped me refine the concept of this research, and his deep interest in this topic and unfailing encouragement are highly appreciated.

## Authors’ Information

**This information is presented on the title page**.

